# Mitochondrial Dysfunction and Neuronal Anomalies in *POLG* Mutant Midbrain Organoids

**DOI:** 10.1101/2023.09.27.559684

**Authors:** Anbin Chen, Tsering Yangzom, Gareth John Sullivan, Kristina Xiao Liang

**Affiliations:** Department of Neurosurgery, Xinhua Hospital, Shanghai Jiaotong University School of Medicine, Shanghai, China; Center for Diagnosis and Treatment of Cranial Nerve Diseases, Shanghai Jiao Tong University, Shanghai, China; Department of Clinical Medicine (K1), University of Bergen, Bergen, Norway; Centre for International Health, University of Bergen, Bergen, Norway; Department of Molecular Medicine, Institute of Basic Medical Sciences, University of Oslo, Oslo, Norway; Institute of Immunology, Oslo University Hospital, Oslo, Norway; Hybrid Technology Hub Centre of Excellence, Institute of Basic Medical Sciences, University of Oslo, Oslo, Norway; Department of Pediatric Research, Oslo University Hospital, Oslo, Norway

**Keywords:** Midbrain organoids, Pluripotent stem cells, Dopaminergic neurons, Mitochondrial dysfunction

## Abstract

Human pluripotent stem cell-derived midbrain organoids offer transformative potential for elucidating brain development, disease representation, and therapeutic innovations. We introduce a novel methodology to generate midbrain-specific organoids from both embryonic stem cells (ESCs) and induced pluripotent stem cells (iPSCs). By employing tailored differentiation cues, notably dual-SMAD inhibition combined with FGF-8b and Sonic hedgehog agonist purmorphamine, we direct pluripotent stem cells towards a midbrain lineage. These organoids, growing beyond 2mm in diameter, house diverse neuroepithelial cell populations. Their midbrain character is affirmed by the pronounced expression of midbrain-specific markers and the absence of forebrain and hindbrain indicators. Critically, these organoids differentiate into dopaminergic neurons characteristic of the midbrain, displaying both morphological sophistication and electrophysiological vigor. Additionally, our experiments with POLG iPSC-derived midbrain organoids revealed a marked loss of dopaminergic neurons and diminished expression of genes governing mitochondrial pathways. This evidence underscores the model’s potential in simulating mitochondrial diseases and neurodegenerative conditions, notably Parkinson’s disease. Our protocol thus emerges as a pivotal instrument for crafting functionally adept, midbrain-centric organoids, paving avenues for advanced studies in midbrain evolution, disorders like Parkinson’s disease, and their interplay with mitochondrial dysfunction.

## Introduction

The human brain, characterized by its intricate structural, cellular, and molecular patterns, continues to stand as a formidable challenge in modern research. Unlocking its vast mysteries, especially in the domains of neuroscience and disease modeling, necessitates in vitro models that accurately capture the brain’s profound complexity. The scientific journey over the past years has been marked by notable strides in manipulating the differentiation of human pluripotent stem cells (hPSCs) into region-specific neuronal subtypes, providing unparalleled insights into human brain development, disease modeling, and therapeutic interventions [1-4].

However, while indispensable, the traditional two-dimensional neuronal cultures derived from hPSCs often miss capturing the full spectrum of cellular composition and structural richness of the human brain [5, 6]. This gap led to the rise of brain organoid technology-a cutting-edge in vitro paradigm emulating the sophisticated architecture and functions of the human brain [7, 8]. These pioneering methodologies have harnessed the self-organizing propensities of PSCs, guiding them to mimic the early milestones of brain development [9-11].

As we plunge deeper into the understanding of the human brain, the intricate neuronal networks stand out, highlighting the importance of three-dimensional (3D) models. These models, more than ever, are essential in reproducing these networks, especially when the spotlight is on neurodegenerative disorders [12, 13]. Facilitating a more ’natural’ milieu for neuronal evolution, 3D models present a closer-to-life representation, both in terms of morphology and physiological dynamics [14, 15]. Indeed, the past decade has been transformative. A multitude of 3D brain models have surfaced, each aiming to emulate specific facets of the human brain’s spatial and functional attributes. These models, ranging from all-encompassing brain organoids to those targeting individual brain sectors, have unveiled groundbreaking insights into conditions like microcephaly, Batten disease, and notably, Parkinson’s disease (PD)[10, 16, 17].

PD, a progressive neurodegenerative disorder, primarily affects the midbrain, leading to a gradual decline in motor functions. A deeper understanding of disease etiology and mechanisms has been a longstanding quest. Concurrently, POLG disease, a disorder stemming from mutations in the DNA polymerase gamma, often manifests with symptoms resembling PD [18]. This similarity suggests shared pathways and mechanisms, underscoring the potential of a unified model for study.

In this context, we’ve pioneered 3D human midbrain organoids (MOs) springing from iPSCs. These 3D marvels promise to illuminate the shadowy corridors of POLG pathology. By crafting and delving into these midbrain-centric organoids, we’re charting a course that reshapes our understanding of neurodevelopmental and neurodegenerative phenomena. This evolution underscores the transformative potential of 3D paradigms, poised not merely as scholarly instruments, but as groundbreaking linchpins in therapeutic development and precision medicine, heralding novel vistas in PD and POLG disease mitigation.

## Methods

### Ethics Approval

The Western Norway Committee for Ethics in Health Research approved this study (REK nr. 2012/919). All participating patients provided written informed consent, and the research conformed to the principles laid out in the WMA Declaration of Helsinki and the Department of Health and Human Services Belmont Report.

### Culture of iPSCs

We generated patient iPSCs from parental fibroblasts carrying *POLG* mutations, one homozygous for c.2243G>C; p.W748S (WS5A) using previously described methods [19-22]. Fibroblast lines Detroit 551 and CCD-1079Sk were reprogrammed as disease-free controls iPSCs from Detroit 551 and CCD-1079Sk fibroblasts [19, 23, 24]. We also used human embryonic stem cell lines (ESCs) line 360 obtained from the Karolinska Institute, Sweden [25] as an internal control for hMO differentiation.

The iPSCs and ESCs were cultured on Geltrex (Thermo Fisher Scientific, #A1413302) -coated 6- well plates (Thermo Fisher Scientific, #140675) using Essential 8 Basal medium (Thermo Fisher Scientific, #A1516901) supplemented with a Y-27632 dihydrochloride Rock Inhibitor (Biotechne Tocris, #1254) for the first 24 hours post-plating. The culture medium was refreshed daily, and iPSCs (about 70%-80% confluent) were split using 0.5mM EDTA (Thermo Fisher Scientific, #15575038). We regularly monitored all cell cultures for mycoplasma contamination using the MycoAlert™ Mycoplasma Detection Kit (Lonza, #LT07-218).

### Generating and Culturing MOs

To manipulate the specification of neural progenitor cells, we split iPSCs at 70% confluence and seeded them onto Geltrex-coated plates in E8 medium. After 24 hours, we switched to a neural induction medium supplemented with 10 µM SB431542 (Tocris Bioscience, #1614), 10 µM N-Acetyl L cysteine (NAC, Sigma-Aldrich, #A7250) and 2 µM AMPK inhibitor Compound C (EMD Millipore, #US1171261 1MG) in Chemically Defined Medium (CDM) containing 50% Iscove’s Modified Dulbecco’s Medium (IMDM, Thermo Fisher Scientific, #21980 065), 50% F12 Nutrient Mixture (Ham) liquid with GlutaMAX™, 5 mg/ml bovine serum albumin (BSA) Fraction V (Europa bioproducts ITD, #EQBAC62 1000), 1% (v/v) Lipid 100 X (Thermo Fisher Scientific, #11905 031), 450 μM 1 hioglycerol (Sigma-Aldrich, #M6145 25ML), 7 μg/ml insulin (Roche, #11376497001), and 15 mg/ml transferrin (Roche, #10652202001). Once cells reached the neural epithelial stage at day 5, we detached them using collagenase IV and then transferred them into non-treated culture dishes. From day 5 onwards, we maintained the generated neurospheres inCDM supplemented with FGF-8 (R&D Systems, #423-F8) for 7 days. From day 12, we switched to a medium containing Purmorphamine (PM, EMD Millipore, #540220-5MG) and FGF-8 for final patterning towards midbrain floor plate precursors. From day 19 onward, we embedded the MOs in Matrigel Matrix and added Brain-derived neurotrophic factor (BDNF, PeproTech, # 450- 02) and Glial cell line-derived neurotrophic factor (GDNF, PeproTech, #450-10) to the culture medium for final maturation up to four months.

### Midbrain Dopaminergic (mDA) Neuron Differentiation

For DA neuronal differentiation, MOs generated from iPSCs were maintained in CDM supplemented with 100 ng/ml FGF8b for 7 days to initiate mDA progenitor cell induction. For the next 7 days, the medium was changed to CDM supplemented with 1 μM PM and 100 ng/ml FGF-8. Suspension cultures were terminated by dissociation of spheroids into single cells with TrypLE™ Express (Thermo Fisher Scientific, #12604039), followed by triturating and subsequent plating in monolayer. mDA neurons were matured in DA medium: Chemically defined medium (CDM) editing plates supplemented with 10 ng/ml BDNF and 10 ng/ml GDNF on poly-L-ornithine (Sigma-Aldrich, #P4957) and laminin (Sigma-Aldrich, #L2020) coated plates.

### Immunostaining of iPSCs and mDA Neurons

Cells designated for this process, including iPSCs and mDA neurons, were first fixed in a 4% paraformaldehyde solution (PFA, VWR, #100503 917). They were then prepped for staining by blocking in 1× PBS, 10% normal goat serum (Sigma-Aldrich, #G9023), and 0.3% Triton™ X-100 (Sigma-Aldrich, #X100-100ML). Overnight incubation at 4°C with primary antibodies followed, targeting specific markers for iPSCs with anti-SOX2, anti-OCT4, anti-SSEA4, and anti-Nanog, and for mDA neurons with anti-TH, anti-Tuj1, anti-MAP2, anti-Synaptophysin, and anti-PSD95. Next, we rinsed in PBS for 15 minutes, then the samples were exposed to secondary antibodies for an hour at room temperature, in a humid and light-protected environment. The slides were then mounted using Fluoromount-G supplemented with DAPI (Southern Biotech, #0100-20).

### Staining Procedure for MOs

For MO cells, the initial step was spreading them onto cover slips, allowing them to dry at room temperature, followed by fixation in 4% PFA. The spheres were washed twice in PBS, before being covered in a 20% sucrose PBS solution, secured, and stored overnight at 4°C. The next day they were blocked for 2 hours at room temperature and then primary antibodies were added and incubated overnight at 4°C. The next day they were washed for 3hours with regular PBS buffer exchanges. Then secondary antibody was added and incubated overnight at 4°C in a humid, dark setting. The final step was mounting the coverslips using Fluoromount-G supplemented with DAPI. A detailed list of antibodies used is in Table S1.

### Microscopy and Image Quantification

We capture images using a Leica TCS SP8 confocal laser scanning microscope. The image acquisition software used was Leica LAS X, and Fiji (Fiji Is Just ImageJ) was used for image processing and analysis. Randomly selected regions (6-10) within the cortical layer for quantitative evaluation. To measure fluorescence intensity, we converted single-channel images to 8-bit images using Image J. To eliminate errors caused by manually selecting thresholds for different photos, we used default thresholds. Next, we choose the default algorithm and parameters. Finally, the fluorescence intensity was measured by the mean gray value (Mean), and the calculation formula is: Mean = integral density (IntDen)/area. Data were analyzed and graphed using GraphPad Prism 8.0.2 software (GraphPad Software, Inc).

### Flow Cytometry Analysis

The cells were first detached using TrypLE™ Express (Thermo Fisher Scientific, #12604013) and subsequently fixed in 1.6% PFA at room temperature for a duration of 10 minutes. Following fixation, they underwent permeabilization using 90% methanol kept at −20°C for 20 minutes. A blocking step was performed using a solution of 0.3 M glycine, 5% goat serum, and 1% BSA in PBS. Subsequently, cells were stained using primary antibodies targeting TH and DAT, followed by an incubation with a diluted secondary antibody (1:400). The stained cells were immediately analyzed on a BD Accuri™ C6 flow cytometer. Data obtained was processed using the Accuri™ C6 software, utilizing dot plots of SSC-H/SSC-A and FSC-H/FSC-A to omit duplicate events. For precision, over 40,000 events were documented for each sample. A detailed list of the antibodies used is in Table S1.

### Single Cell RNA Sequencing (scRNA-seq) and Data Analysis

### Organoid Dissociation and Single Cell Isolation

Organoids were removed from the medium and washed oncein PBS. The organoids were then finely minced into 1-2 mm pieces using ophthalmic scissors. Next, the organoid fragments were digested in 2 ml of CellLive™ Tissue Dissociation Solution (Singleron Biotechnologies, #1190062) in 15 ml centrifuge tubes (Sarstedt, #62.5544.003) at 37°C for 15 minutes with continuous agitation on a heated shaker. The organoids were periodically checked by light microscopy for dissociation. After digestion, the dissociated organoids were passedthrough a 40 µm sterile filter (Greiner, #542040). Cells were then centrifuged at 350 x g for 5 minutes at 4°C and the resulting cell pellet was resuspended in 1 ml PBS. To assess cell viability and numbers, cells were stained with 0.4% w/v trypan blue solution (Gibco, #15250-061). Cell number and viability were determined using a hemocytometer under a light microscope.

### Data Preprocessing

Pre-processing of Fastq data was conducted using CeleScope® (v.1.14.1; www.github.com/singleron-RD/CeleScope; Singleron Biotechnologies GmbH) to generate raw data, using default parameters. Low quality reads were removed. Sequences were mapped using STAR (https://github.com/alexdobin/STAR) and the human reference GRCh38 and genes were annotated using Ensembl 92. The reads were assigned to genes using featureCount (https://subread.sourceforge.net/) and the cell calling was performed by fitting a negative bimodal distribution and determining the threshold between empty wells and cell-associated wells to generate a count matrix file containing the number of Unique Molecular Identifier (UMI) for each gene within each cell. Downstream analysis was done in scanpy package [26] implemented in Python software and Seurat library [27] in R software. QC metrics, such as number of genes detected per cell (nFeature_RNA) and percentage of mitochondrial UMI (percent_mt) were extracted from the gene count matrix. To remove non-viable cells, cells with high percentage of mitochondrial counts (>20%) were filtered out. To avoid doublets, the cells with a high number of detected genes (>6500) and high number of detected UMI (>30000) were removed. Cell debris characterized with a low number of detected genes (<200) were also removed from the data.

### Clustering and Cell Type Annotations

Data integration was conducted using Harmony (https://github.com/immunogenomics/harmony) [28] via scanpy pipeline. A clustering analysis was conducted from the integrated 4 samples (control 1, control 2, disease and treated) using Scanpy. On the combined dataset, we computed the neighbourhood graph using the function scanpy.pp.neighbors with the option of n_neighbors = 10 and n_pcs = 10. We conducted unsupervised clustering of cells applying the leiden algorithm using the function scanpy.tl.leiden with resolution 1.

Clusters were annotated based on the expression of known markers identified from several literatures. DA progenitors were annotated for their highly expressed markers such as NEUROG2, NHLH1 and SOX4 [29, 30]. DA-0 was annotated based on the high expression of ErbB4 known to be expressed in DA neurons and low level of SYT1 a post-synaptic marker[31]. DA-1/2 was annotated since it expressed a well-known DA neuronal marker NR4A2 as well as DCX and SYT1 among others [32, 33]. Meningeal cells were annotated based on the high expression of COL1A1, COL1A2, and LUM. Neural progenitors were annotated based on the expression of TOP2A, MKI67, and HMGB2. Cluster 2, 5, and 14 were annotated as Oligodendrocytes based on expression of RBFOX1 [34]. Astrocytes was annotated based on high expression of GFAP, AQP4, EDNRB and RFX4. Oligodendrocyte progenitors were annotated by the high-level expression of OLIG2. Cluster 7 was annotated as Radial glial cells which marked by high expression of FABP7. Glial progenitor was annotated since they expressed TOP2A, MKI67, and HMGB2 as well as FABP7 [32, 35]. Ventral midbrain neurons (VMN) had gene markers that are close to but not identical as DA neurons and annotated based on celltypist [36] using developing human brain models [37] to find the closest cell type using logistic regression models to be ventral midbrain neurons. The gene expression for the cell cluster annotation was listed in Figure 5C. Control 1 (Detroit 551) and control 2 (CCD-1079Sk) were combined to make a control sample with further downstream analysis. Clustering and cell type annotations were kept the same with the cells for the control samples.

### Differential Expression and Pathway Analysis

Differential expression analysis was performed using scanpy.tl.rank_genes_groups on the identified cell clusters and between different cell types. The sample-based cell proportions were calculated as well as the absolute counts for individual clusters in the subgroup. Clusters were custom grouped, and differential expression analysis was performed using Wilcoxon rank sum test with benjamini-hochberg p-value correction methods.

### Enrichment Analysis

For the enrichment analysis of Gene Ontology (GO) and the Kyoto Encyclopedia of Genes and Genomes (KEGG), genes marked as upregulated and downregulated were meticulously analyzed. An online tool, available at https://www.bioinformatics.com.cn, was leveraged to effectively conduct both the GO and KEGG enrichment analyses, as described previously [38]. This comprehensive analysis aids in the identification and understanding of significant genes and pathway enrichments, contributing to the broader perception of their roles and interactions in the biological context.

### Statistical Analysis

Our data representation takes the form of mean ± standard deviation (SD), with a minimum sample size of three. We assessed the normality of data distribution using the Shapiro-Wilk test and detected any outliers using the ROUT method. When dealing with variables of non-normal distribution, we determined statistical significance using the Mann-Whitney U-test. In contrast, variables with normal distribution underwent analysis using a two-sided Student’s t-test. All graph generation and statistical analyses were conducted using GraphPad Prism 8.0.2 software. We deemed a p-value of ≤ 0.05 to indicate statistical significance.

## Results

### Generation of 3D MOs

We generated midbrain organoids (MOs) from pluripotent stem cells that displayed markers like NANOG, OCT4, and SOX2, as depicted in Figure 1A. Our differentiation method, illustrated in Figure 1B, involved a dual-SMAD technique complemented by the inclusion of FGF-8b and the Sonic hedgehog (SHH) stimulant PM. To guide the neuroectoderm towards the floor plate, we initiated the differentiation of stem cells into neural rosettes via dual SMAD inhibition and the introduction of N-Acetylcysteine (NAC). By dissociating these cells and cultivating them in a suspension environment, we were able to form 3D neurospheres. These were consistently agitated on an orbital shaker at 80 rpm. Subsequent treatments involved caudalization using FGF-8b for a week, followed by a ventralizing procedure with a blend of FGF-8b and PM for another week. The MOs that resulted were then encapsulated in Matrigel droplets for structural reinforcement and were matured further by adding BDNF and GDNF, as shown in Figure 1C. Within a month, these organoids achieved over 2 mm in diameter and housed numerous neuroepithelial cells, as visualized in Figures 1D and 2B. For our experiments, we utilized one ESC variant (ESCs 360) and two distinct human iPSC models (Detroit 551 and CCD-1079Sk lines).

**Figure 1.**
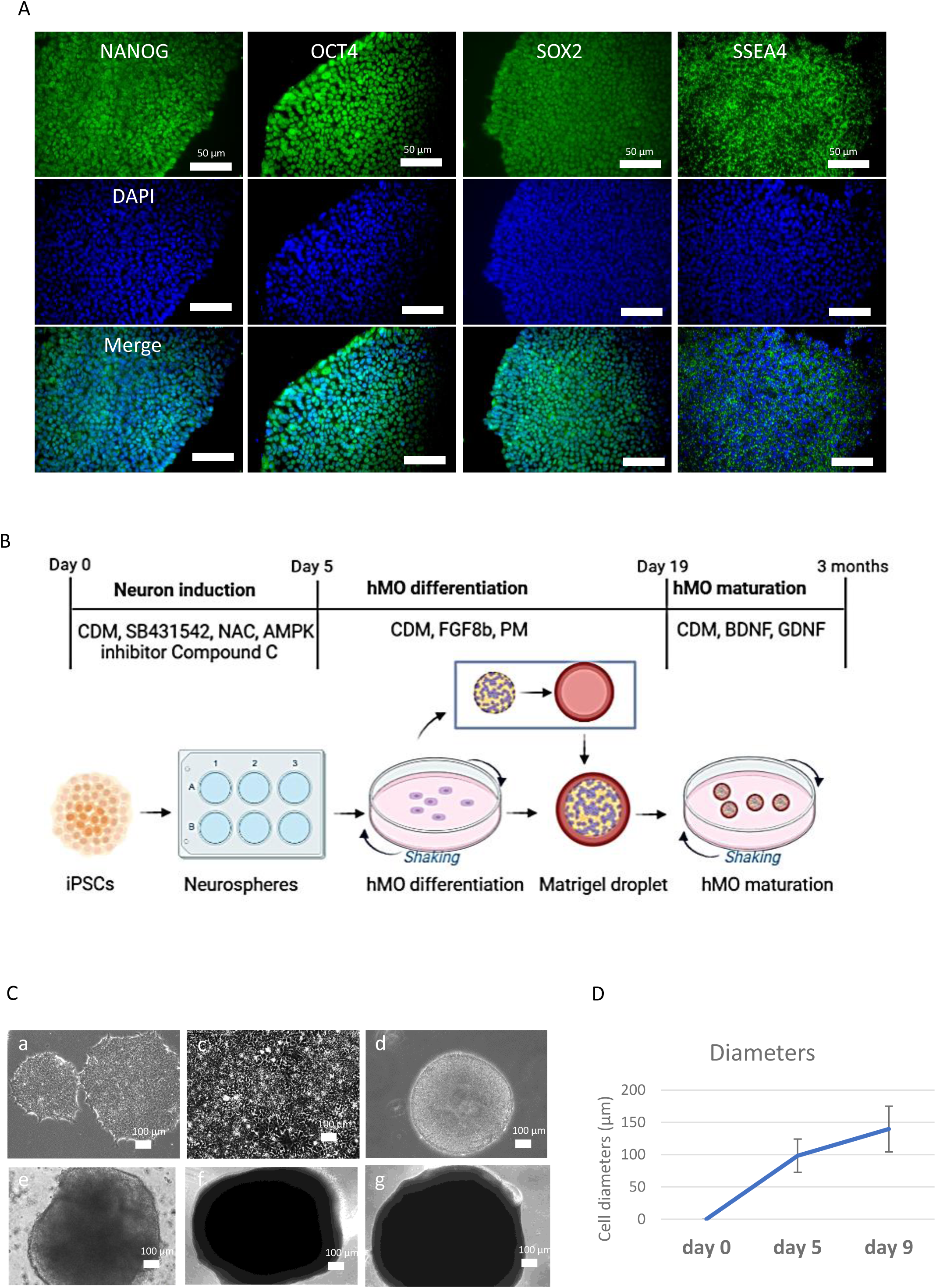
**Generation of MOs from iPSCs and ESCs.** A. Representative fluorescent images of control iPSCs labeled with Nanog, OCT4, SOX2, and SSEA4. Nuclei were stained with DAPI (blue). Scale bar is 50 µm. B. Schematic representation of control iPSC or ESC differentiation into MOs. C. Representative phase-contrast images of iPSCs (a), neuroepithelium (b), DA progenitors (c) and MOs generated from control iPSCs (d) and ESCs (e) on days 19 and 20 (d and e). Scale bar is 100 µm. D. Quantification of cell diameter during differentiation

**Figure 2.**
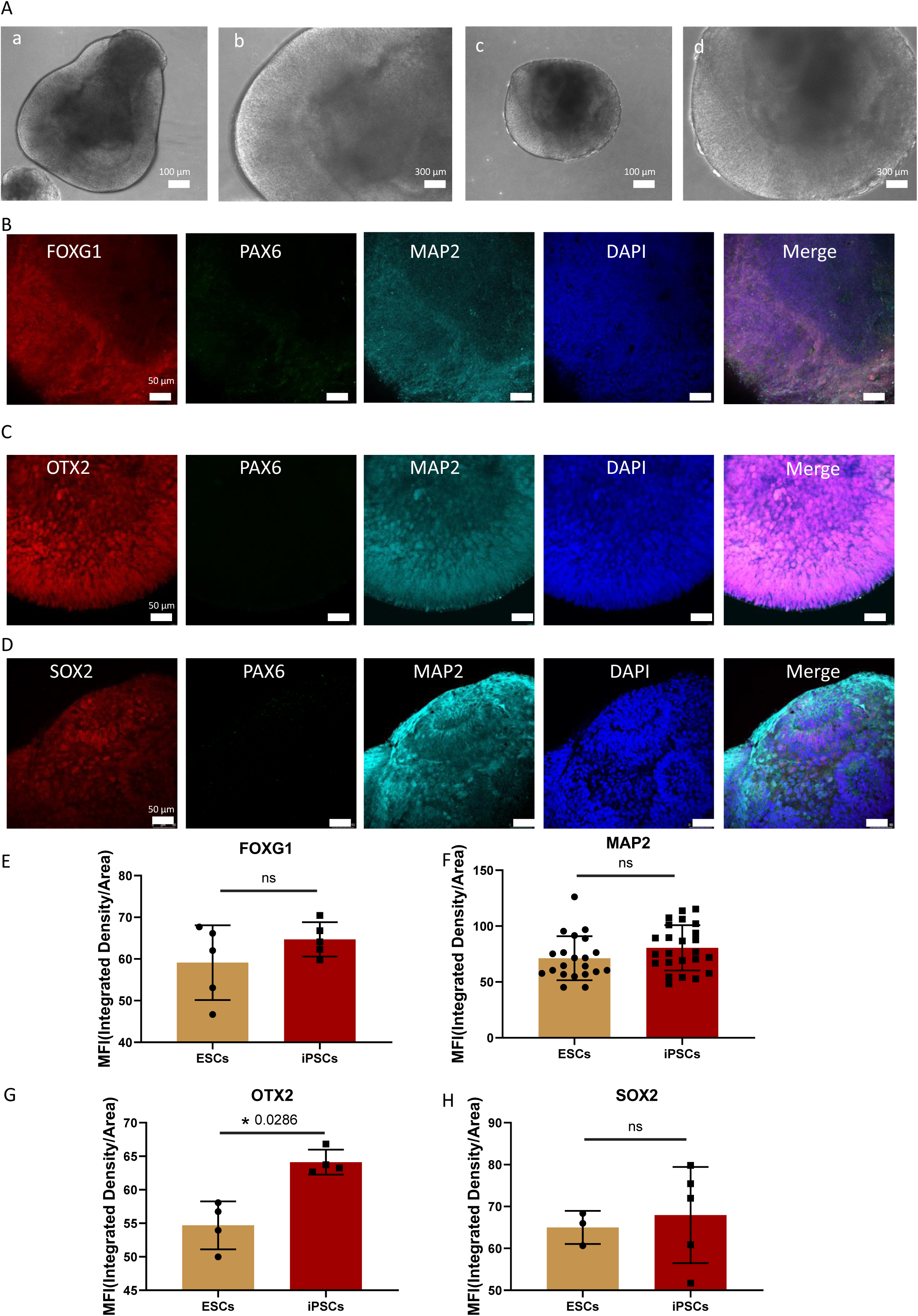
**Characterization of markers of midbrain progenitor cells.** A. Phase-contrast images of 3-month-old MOs from ESCs (a: 10x magnification and b: 20x magnification) and control iPSCs (c: 10x magnification and d: 20x magnification). Scale bar is 100 µm. B-D. Fluorescent staining for markers of progenitor cells in the MOs at 3 months of age, including triple staining for FOXG1/PAX6/MAP2 (B), OTX2/PAX6/MAP2 (C), OTX2/PAX6/MAP2 (D). Nuclei were stained with DAPI (blue). Scale bar is 50 µm. E-H: Quantification of immunofluorescence staining in B-D.

In essence, we successfully crafted midbrain organoids from pluripotent stem cells, marked by NANOG, OCT4, and SOX2. Utilizing a dual-SMAD strategy enhanced with specific treatments like FGF-8b and PM, we guided their differentiation to achieve a distinct midbrain identity. The resulting organoids, maturing to over 2 mm in a month, highlight the efficacy of our protocol in fostering robust neuroepithelial cell growth, especially when considering the consistent outcomes across diverse stem cell lines, including one ESC and two iPSC variants.

### Characterization of Midbrain-Specific Markers

On the 60th day, we examined MOs for the presence of the transcription factor PAX6 to distinguish between midbrain and forebrain identities. Interestingly, the absence of PAX6 expression confirmed the midbrain character (Figures 2B-D). To further confirm a forebrain lineage, we checked for the transcription factor FOXG1, known for its critical role in forebrain specification, and found its presence (Figure 2B). Additionally, MOs displayed positive SOX2 staining, a hallmark of neural progenitor cells, along with the mature neuron marker MAP2, signifying the formation of neurons at this juncture (Figures 2B-D). Notably, both ESC- and iPSC-derived midbrain organoids showcased comparable expressions of FOXG1, SOX2, and MAP2 when analyzed for fluorescence intensity (Figure 2B-D).

Delving deeper into the midbrain neuroepithelium’s characterization, OTX2-a transcription factor vital for the mid-hindbrain organizer-manifested within the apical region extending midway through the neuroepithelium (Figure 2D). To pinpoint mDA neurons specifically, we stained for tyrosine hydroxylase (TH), a definitive mDA neuron marker. Notably, cells co-expressing TH and MAP2 were identified, particularly on the neuroepithelium’s apical surface, in MOs at the 60-day mark (Figure 3B). By day 90, these MOs prominently displayed TH co-expression with the neuronal marker class III β-tubulin (TUJ1) (Figure 3C). Fluorescence intensity measurements further demonstrated that OTX2 (Figure 3D) and TH (Figure 3E) expressions in iPSC-derived MOs paralleled those from ESC-derived samples.

**Figure 3.**
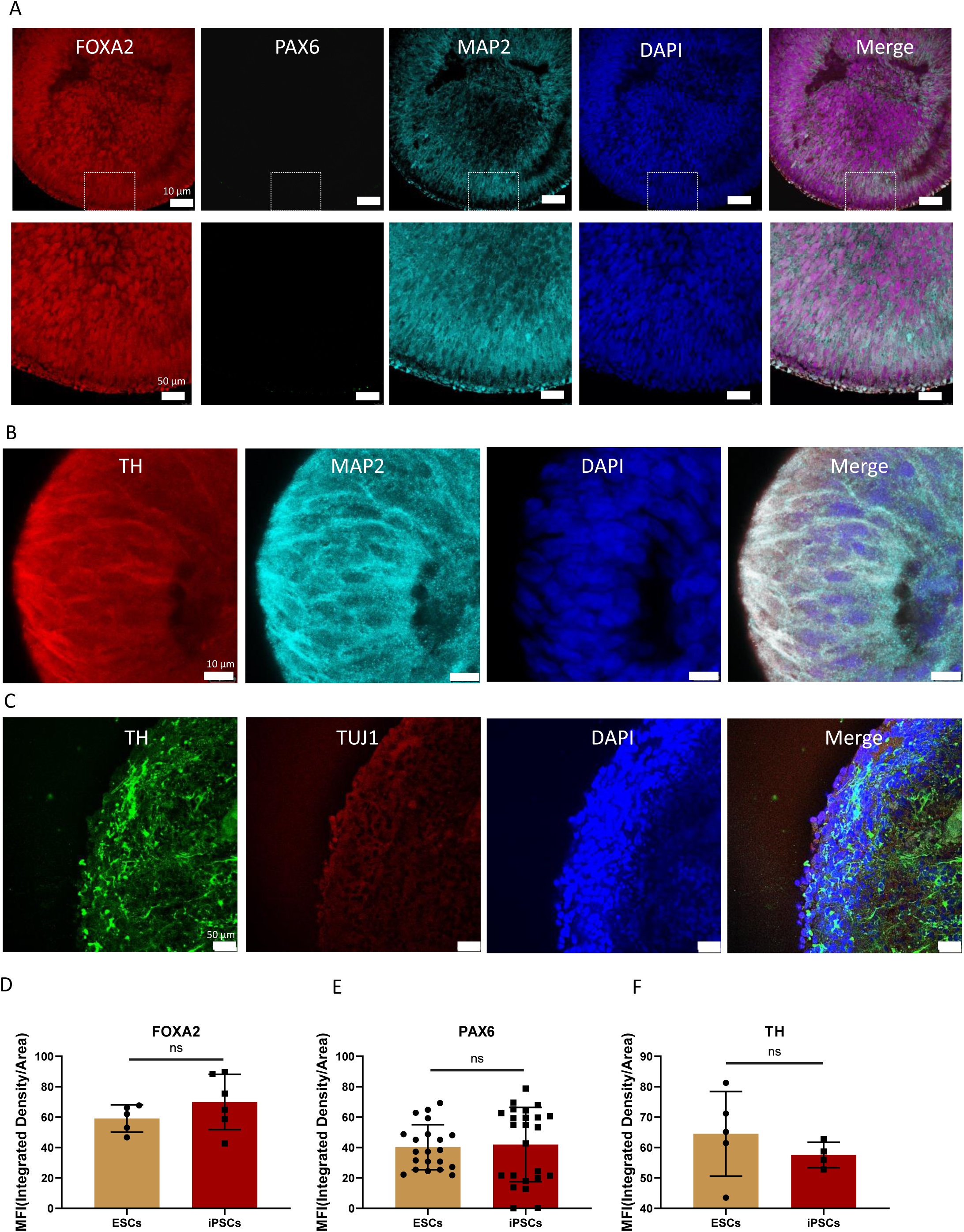
**Characterization of midbrain markers in MOs.** A-C. Fluorescent staining for markers of progenitor cells in the MOs at 3 months of age, including triple staining for FOXA2/PAX6/MAP2 (A), TH/MAP2 (B), TH/TUJ1 (C). Nuclei were stained with DAPI (blue). Scale bar is 50 µm. D-F. Quantification of immunofluorescence staining for FOXA2, PAX6 and TH in A-C.

In summary, 60-day matured MOs distinctly showcased midbrain characteristics, evident from the absence of PAX6 and the presence of FOXG1. Neural progenitor markers, coupled with mature neuron indicators, highlighted the advanced neuronal development within these organoids. Both ESC- and iPSC-derived MOs presented with remarkably similar expression patterns, emphasizing the consistency in neuronal development across derivation methods.

### Neuronal Differentiation of ESC/iPSC-Derived MOs

Following the assessment of the neuronal differentiation potential of MOs, we probed their capability to mature into mDA neurons. Upon dissociating 2-month-old MOs to singular cells and nurturing them for another month, we noticed the emergence of intricate neuronal networks (Figure 4A and B). Flow cytometry analysis illuminated that a vast majority of these neurons were TH+ (98.2%, Figure 4B). Furthermore, a significant fraction of these neurons exhibited dopamine transporter (DAT) expression, accounting for 76.6% (Figure 4B and C). Delving deeper with immunostaining targeting a selection of mDA neural lineage markers, it was evident that the mDA neurons derived from the midbrain organoid co-expressed DAT, TH, and MAP2. This was notably pronounced in the matured neurons (Figure 4C), underscoring a robust and efficient differentiation into mDA neurons from both ESC and iPSC sourced MOs.

**Figure 4.**
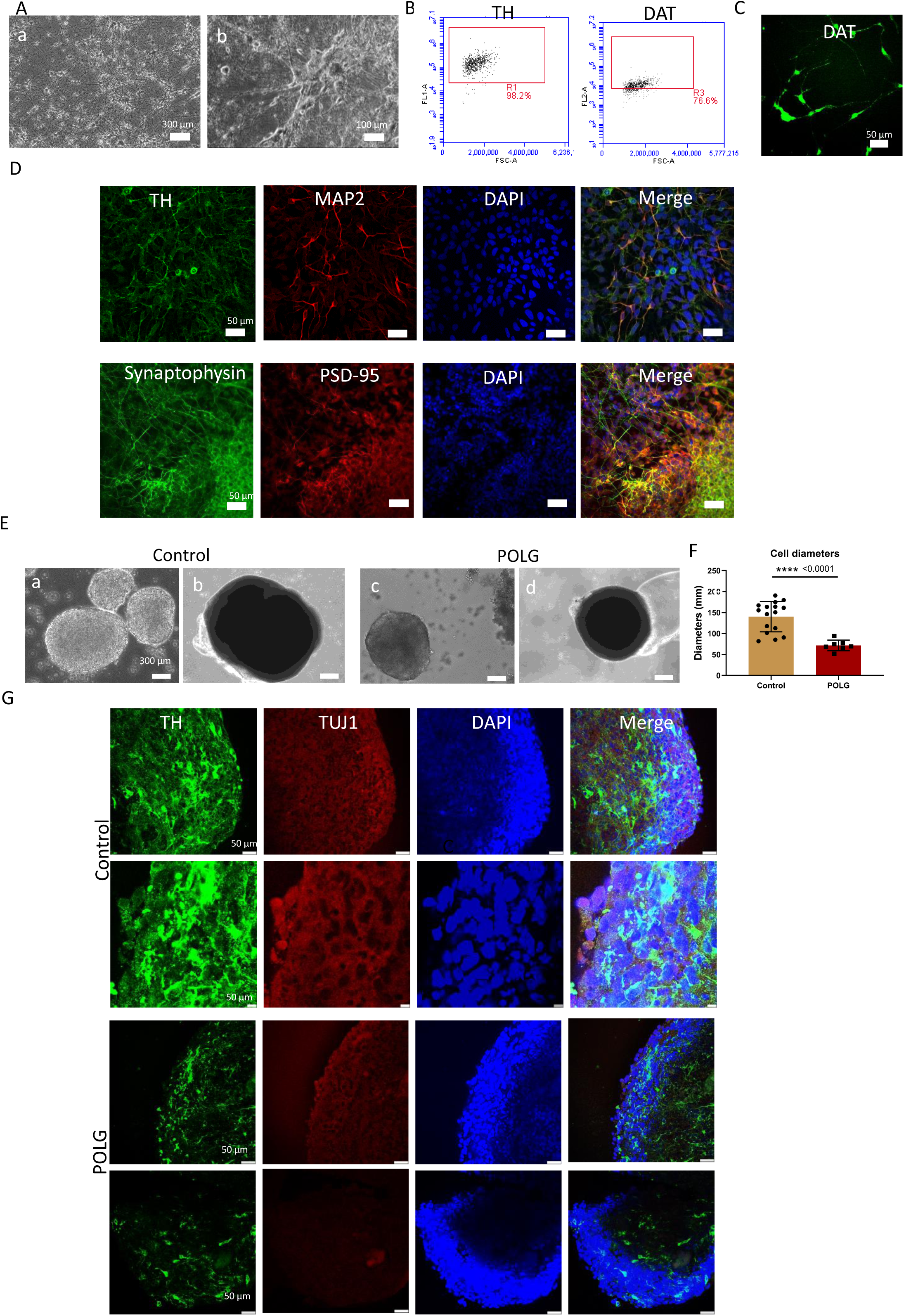
**Differentiation and characterization of mDA neurons from POLG patient and control iPSCs and generation of MO.** A. Phase contrast images of mDA neurons from ESCs (a) and control iPSCs (b). Scale bar is 100 µm. B. Flow cytometric analysis of the mDA neural markers TH and DAT in MO-generated mDA neurons. C. Fluorescent staining of the mDA neural marker DAT in mDA neurons. Nuclei were stained with DAPI (blue). Scale bar is 50 µm. D. Fluorescent staining of mDA neurons for the mDA neural marker TH, the mature neural marker MAP2, the presynaptic marker synaptophysin, and the postsynaptic marker PSD95 in mDA neurons. Nuclei were stained with DAPI (blue). Scale bar is 100 µm. E. Phase contrast images of MO generated from control and POLG patient iPSCs. Scale bar is 100 µm. F. Comparison of cell diameter quantification in control and POLG MO.

At the close of a 4-week maturation period, our attention turned to further markers characterizing mDA neurons. In a bid to visualize neurofilaments, we utilized MAP2 staining and concurrently probed synaptic markers. The mDA neurons were stained for synaptophysin, a principal synaptic vesicle protein, p38 (serving as a presynaptic marker), and synapse-associated protein 90 (referred to as PSD-95 or SAP-90) – a protein integral for anchoring receptors and other signaling molecules (playing a postsynaptic role). Remarkably, our mDA neurons manifested positive co-expression of synaptophysin and PSD-95. The observed pattern suggests these neurons might be forming synaptic connections (Figure 4D).

In summary, our MOs exhibited a robust capability to mature into mDA neurons, with intricate neuronal networks forming over time. A significant majority of these neurons expressed key DA lineage markers, underlining an efficient differentiation process from both ESC and iPSC sources. The maturation culminated with these neurons showcasing signs of synaptic connections, emphasizing their functional potential and structural integrity.

### POLG Midbrain Organoids Exhibited a Reduction in mDA Levels and Displayed Downregulated Single-Cell Transcriptomic Profiles

Next, we applied our protocol to study the mitochondrial disease with *POLG* mutation. We generated patient specific iPSC derived POLG MOs. The resulting POLG MOs exhibited irregular morphology (Figure 4E). Additionally, the POLG MOs were significantly smaller in size as compared to control MOs (Figure 4F). We next assessed TH and TUJ1 marker staining, we observed a reduction in their expression in POLG MOs as compared to controls (Figure 4G).

To delve deeper we investigated the transcriptomic profiles at the single-cell level, by single cell RNA sequencing (scRNA-seq) analysis. For comparison purposes, we combined Detroit 551 and CCD-1079Sk lines to serve as a control against the patient-derived organoids (Figure 4F). Analysis from scRNA-seq indicated that both control (in total 17698 cells) and patient-derived MOs (in total of 4871 cells) consisted of DA neurons, Oligodendrocytes, Radial glial cells, Oligodendrocyte progenitors, Glial progenitors, Ventral midbrain neurons, DA-0 cells, Astrocytes, Neural progenitors, DA progenitors, and Meningeal cells, identified based on their gene expression patterns (Figure 5A-D).

In our examination of POLG patient-derived MOs, there was a marked decrease in DA neurons. These comprised a mere 15.25% (from a total of 743 cells) as opposed to the 30.2% (from a comprehensive 5,353 cells) observed in control Mos (Figure 5D). Moving on to gene expression, the patient MOs demonstrated a significant deviation from controls. There was a conspicuous downregulation (Figure 5E). A detailed breakdown revealed 543 DEGs being upregulated, whereas a significant 2,915 DEGs were downregulated.

To delve deeper into these variances, we utilized GO enrichment analyses to decipher the probable biological processes being affected. We found the most affected processes in the downregulated DEGs (Figure 5G). These prominently include regulation of trans-synaptic signaling, modulation of synaptic transmission, and synapse organization. The data illuminates a considerable disruption in synaptic activities, especially emphasizing the synaptic vesicle cycle. Furthermore, indications towards anomalies in axonal development, axonogenesis, and potential dysregulations in neuronal communication, particularly in the membrane potential and neurotransmitter level adjustments, were discerned. Emphasizing the role of synaptic mechanisms, our findings suggest significant perturbations in synaptic signaling, organization, and transport processes. Given the context of established markers like mitochondrial protein complexes, it is evident that fundamental cellular systems are perturbed. Moreover, an emapplot analysis of the downregulated DEGs in POLG MOs, as compared to controls, shed light on the pivotal "modulation of chemical synaptic transmission." This process appeared to be at the nexus of several affected biological processes, a relationship graphically illustrated in Figure S1."

In our exploration of GO cellular components (Figure 5H), we identified key areas of concern. Foremost among these were components linked with synaptic structures and functions: the synaptic membrane, postsynaptic membrane, glutamatergic synapse, and neuron-to-neuron synapse. This significant focus on synaptic entities points to a potential disturbance in synaptic interactions and overall neuronal communication within POLG MOs. Parallelly, there was a discernible shift in mitochondrial components. The mitochondrial inner membrane and the mitochondrial protein complex emerged prominently in our findings. Such shifts in mitochondrial structures were particularly noteworthy, echoing established insights into their crucial roles in energy metabolism and overall mitochondrial operations. In light of these findings, the data suggests a multifaceted disruption both at the synaptic and mitochondrial levels. A deeper graphical dive into these components (Figure S2) reveals the intricate and pivotal role of the ’synaptic membrane.’ This underscores the importance of these elements not just individually but in their complex, interdependent roles in preserving cellular health and operations."

In our further analysis centered around GO molecular functions (Figure 5G), several critical functions emerged. The data prominently showcased molecular functions related to neural communication, with glutamate receptor binding and ion channel binding taking center stage. Such findings hint at possible disruptions in neurotransmitter activity and the intricate dynamics of ion channels. Moreover, a keen focus on intracellular signaling was evident, with GTPase activity, clathrin binding, and tubulin binding pointing towards a profound impact on cellular dynamics and metabolism. The oxidoreductase activity, particularly involving NAD(P)H and associated compounds, illuminates potential metabolic irregularities. It’s noteworthy to mention the emphasis on amyloid-beta binding, which is congruent with existing insights into neurodegenerative pathways. This could be an indication of vulnerabilities linked to protein misfolding or aggregation. Meanwhile, the enrichment of functions related to purine binding, such as purine ribonucleoside and nucleoside binding, suggests a possible perturbation in nucleotide metabolism or associated signaling processes.

In our study of *POLG* mutation-associated mitochondrial disease using patient-derived iPSCs, distinct abnormalities surfaced. The POLG MOs not only showed altered morphology and size but also reduced expression of essential neuronal markers. Deep scRNA-seq analysis highlighted profound shifts in DA neuron populations and gene expression patterns. Most strikingly, there was a pronounced disruption in synaptic components and mitochondrial structures. Molecular functions critical for neural communication, cellular dynamics, and metabolism were also affected. In essence, POLG MOs exhibited significant synaptic and mitochondrial irregularities, revealing deep-rooted disruptions in neural communication and metabolic processes, underscoring the severity of the disease’s impact at a cellular level.

### POLG MOs Demonstrated Mitochondrial and Neuronal Dysfunctions Through Comprehensive Pathway Analysis

In our comprehensive assessment of downregulated gene profiles in DA neurons derived from patients, two overarching areas of dysfunction emerged: mitochondrial anomalies and neuronal impairments. Mitochondrially, the primary downregulations were evident in GO and MF, focusing on oxidoreductase activities targeting NAD(P)H and quinone compounds. There were also marked deficits in several NADH dehydrogenase activities, ATPase functionalities, and iron-sulfur cluster binding (Figure 6A). Neuronally, there were apparent reductions in activities crucial for neural communication and structural integrity, such as glutamate and GABA receptor binding, synaptic components, and microtubule-associated bindings (Figure 6B).

**Figure 6.**
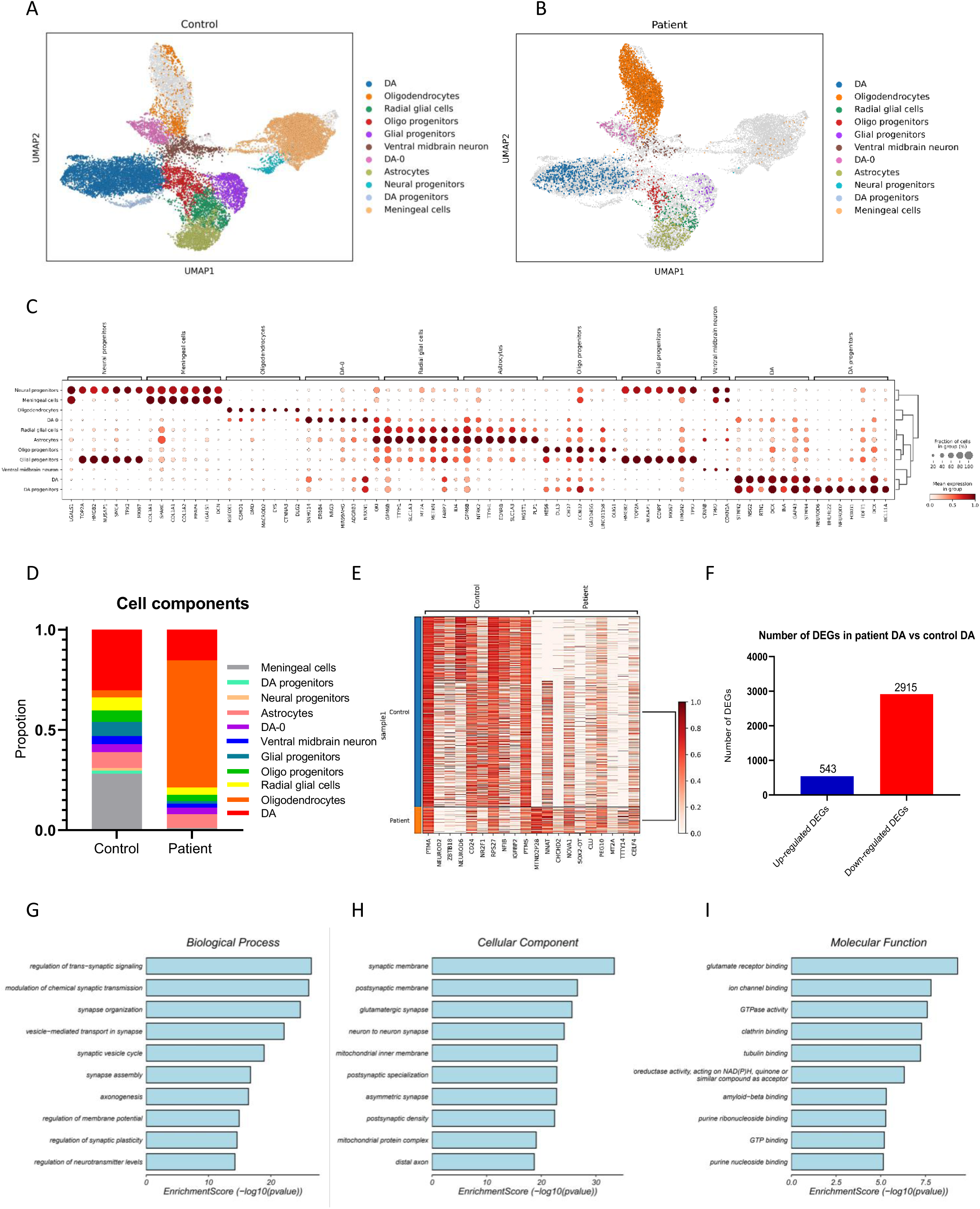
**Sc RNA-seq analysis in control and POLG patient MO.** A, B. UMAP cell annotation analysis of control (A) and POLG patient MO (B) transcriptome expression data. C. Gene expression in different cell clusters in A and B. D. Cell proportions of different cell populations in POLG MOs and controls. E. Heatmap of up-and down-regulated genes in POLG MOs compared to controls. F. Numbers of up-and down-regulated DEGs in MOs in POLG patients compared with controls. G. The top ten GO biological process terms are enriched in downregulated DEGs in POLG in MOs of POLG patients compared with controls. H. The top ten GO cellular component term is enriched in downregulated DEGs in MOs of POLG patients compared with controls. F. The top ten GO molecular functional terms are enriched in downregulated DEGs in MOs of POLG patients compared with controls.

**Figure 7.**
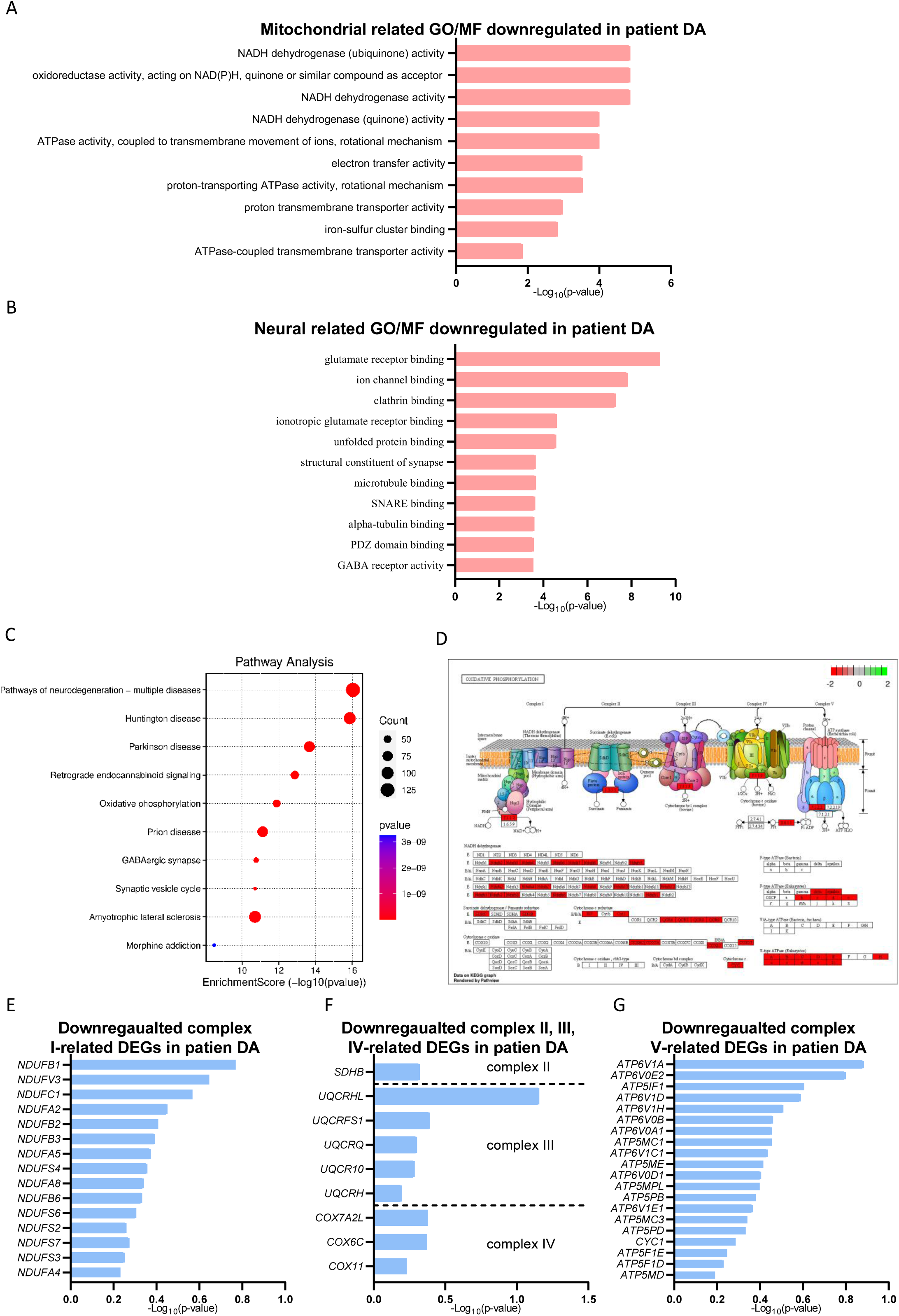
Mitochondrial and neural related GO term and KEGG pathways were downregulated in MOs of POLG patients compared with controls. A. The top ten mitochondria-associated GO molecular functional terms downregulated in MOs of POLG patients compared with controls. B. The top ten neural-associated GO molecular functional terms downregulated in MOs of POLG patients compared with controls. C. The top ten KEGG pathway downregulated in MOs of POLG patients compared with controls. D. Detailed analysis of genes involved in oxidative phosphorylation. E-H. Foldchanges of downregulated DEGs in the mitochondrial respiratory chain complex I-V.

Our KEGG pathway analysis revealed a worrying inclination towards recognized neurodegenerative pathways-Parkinson’s, Huntington’s, and ALS. A standout observation was the pronounced downregulation in the "oxidative phosphorylation" pathway, a critical bioenergetic pathway (Figure S3). Delving deeper, we found a significant clustering of downregulated genes within the mitochondrial respiratory chain complexes, especially complexes I and IV (Figures 6D, 6E-G).

In essence, patient-derived DA neurons exhibited profound mitochondrial dysfunctions and neuronal deficits. The majority of these disparities seem rooted in disruptions to metabolic and bioenergetic processes, painting a concerning picture for the metabolic health and functionality of these neurons in comparison to controls.

### POLG MOs Showed Sympathetically Upregulated Apoptotic and Mitophagy Pathways

Further investigation into the upregulated KEGG pathways in DA neurons within POLG MO highlighted several crucial pathways: Apoptosis, ErbB signaling, Cell cycle, Mitophagy (animal), Regulation of actin cytoskeleton, and the Wnt signaling pathway (Figure S4). Among these, the Apoptosis and Mitophagy pathways stand out, echoing the pivotal role of mitochondrial disturbances in neuronal health. Indeed, apoptosis, driven by mitochondrial dysfunction, has been a focal point in the pathogenesis of several neurodegenerative conditions. Similarly, mitophagy, responsible for the selective degradation of dysfunctional mitochondria, further underscores the delicate balance that exists within neurons to maintain cellular homeostasis.

Moreover, pathways like the ErbB signaling and Wnt signaling, traditionally associated with cellular growth and differentiation, hint at an underlying disruption in neuron maturation and function in the context of *POLG* mutations. The observed changes in the Regulation of actin cytoskeleton emphasize potential disturbances in neuron morphology and synaptic connections.

In summary, DA neurons derived from POLG MO showcased significant mitochondrial anomalies and neuronal irregularities. Central to these perturbations are the disruptions in apoptosis and mitophagy, reflecting the essential role of mitochondria in maintaining neuronal stability. The upregulation of pathways like ErbB and Wnt signaling suggests additional complexities in neuronal maturation and function under *POLG* mutations. Furthermore, alterations in actin cytoskeleton regulation hint at potential challenges in neuronal structure and synaptic integration. Collectively, these findings underscore a compromised metabolic and bioenergetic landscape in these neurons when juxtaposed with their healthy counterparts.

## Discussion

The generation and characterization of 3D MOs from pluripotent stem cells represents an advanced model system to investigate the complexities of human midbrain development, functionality, and associated pathologies. Here, we have demonstrated the meticulous steps required to differentiate pluripotent stem cells into midbrain-specific neurons, mimicking a natural developmental trajectory.

Central to the process of MO generation is the careful use of various differentiation factors. The dual-SMAD approach, complemented by FGF-8b and SHH agonist PM, was successful in differentiating stem cells into neuroectoderm precursors and then onward to midbrain-specific neural cells. This method recapitulates the *in vivo* embryonic neural tube development, and neuroectoderm to the floor plate [39]. Our work supports the role of the FGF-8b and SHH signaling pathways in midbrain patterning, as previously observed [40, 41]. It is notable that these organoids can achieve a diameter of over 2 mm, reflecting the capability of this *in vitro* system to generate MOs of considerable size and complexity.

The use of multiple stem cell lines and the observed concordance of expression patterns of midbrain markers clearly demonstrates the robustness and versatility of our approach in both embryonic and induced pluripotent. Furthermore, the absence of PAX6 expression in our MOs is a positive indication of accurate midbrain specification. PAX6 is pivotal for forebrain development [42]; hence its absence corroborates midbrain identity. However, the presence of FOXG1, SOX2, and OTX2 expression further substantiates a midbrain identity of our organoids. OTX2, in particular, serves as a gatekeeper, distinguishing midbrain from hindbrain territories [43], whereas FOXG1 has been shown to be involved in midbrain patterning [44].

A hallmark achievement of our study is the successful derivation of mDA neurons, as evidenced by TH and MAP2 co-expression. Considering the relevance of these neurons in diseases like PD [45] and POLG disease [1], these organoids offer a promising platform for disease modeling, drug testing, and potential therapeutic applications. The presence of synaptic connections, as indicated by synaptophysin and PSD-95 expression, further demonstrate the maturity and potential functionality of the derived neurons. These synaptic markers have been widely recognized in neurobiology, denoting functional synapses and highlighting the potential of these neurons to integrate into existing neuronal networks [46, 47].

The differentiation of POLG iPSCs into MOs has shed light on potential pathologies stemming from *POLG* mutations. Notably, the irregular and diminished size of POLG MOs, coupled with reduced TH and TUJ1 marker expression, indicates potential disruptions in the development of midbrain-centric neurons. This aligns with prior studies pointing to POLG’s critical role in upholding mitochondrial DNA integrity [48]; its mutations are frequently tied to mitochondrial malfunctions that compromise cellular vitality and operations. From a transcriptomic standpoint, scRNA-seq analysis grants a holistic view of the cellular composition in both patient-derived and control organoids. The striking decline in DA neurons in POLG MOs, nearly 50% less than controls, warrants attention. These neurons are crucial for myriad brain functions, from motor coordination to reward-driven behaviors and several hormonal operations [49-52]. Shortages in these neurons have associations with Parkinson’s disease and other neuromotor ailments [53], hinting that the observed DA neuron deficiency might mirror some clinical manifestations linked to *POLG* mutations. The broad gene downregulation in patient-derived MOs, with almost 3,000 DEGs showing reduced expression, provides a potential glimpse into the molecular disruptions tied to *POLG* mutations. GO enrichment analysis delineates the primary impacted biological activities, cellular elements, and molecular tasks. Remarkably, the notable suppression of pathways connected to synaptic communication and structure points to synaptic malfunction as a possible player in POLG-associated disorders. Such synaptic issues are documented in various neurodegenerative diseases [54], underscoring the pivotal nature of synaptic preservation for neuron survival.

Mitochondrial dysfunction stands out as a shared trait across many neurodegenerative conditions [55]. Notably, the noted changes in the mitochondrial inner membrane and protein complex among the primary affected cellular components resonate with POLG’s established role in safeguarding mitochondrial DNA. The pronounced downregulation of genes connected to mitochondrial elements buttresses the notion that mutations in POLG induce mitochondrial disturbances, subsequently jeopardizing neuronal well-being. Furthermore, the GO molecular function enrichment delineates prospective molecular avenues suitable for therapeutic exploration. The pinpointing of glutamate receptor binding and ion channel binding insinuates potential hiccups in neurotransmission and neuronal excitability. This mirrors research positing excitotoxicity as a potential driver of neuronal demise [55].

The nexus between *POLG* mutations, mitochondrial impairments, and neurodegenerative manifestations has long been underscored (Stumpf and Copeland, 2011). Our comprehensive analysis of patient-sourced DA neurons further solidifies this association, unveiling marked mitochondrial and neuronal irregularities. These insights pave the way for a more refined comprehension of *POLG* mutation repercussions. Our earlier work with 2D systems had hinted at disruptions in DA neurons [20], a notion that the scRNA-seq analysis from patient-derived MOs reaffirms. The remarkable downregulation of genes pivotal to various mitochondrial operations is noteworthy. Especially striking is the pronounced decrease in genes tied to oxidative phosphorylation, a critical cellular energy production pathway. The conspicuous involvement of genes within the mitochondrial respiratory chain complexes accentuates the importance of this pathway in cellular energy dynamics. Given the known criticality of complex I and IV in neuron energy metabolism, the downregulation observed in these gene sets is likely to have deep-seated effects on neuron health [56]. This perspective gains further weight when considering the observed downregulation of genes linked to NADH dehydrogenase activities and ATPase operations. As NADH dehydrogenase-a cornerstone of the mitochondrial respiratory chain-plays a vital role in electron transfer and energy harnessing [57], its dysfunction combined with compromised ATPase activity may culminate in an energy shortfall, thereby jeopardizing neuron functionality and survival.

Within the neuronal framework, the noted shortcomings in glutamate receptor engagement, ion channel associations, and synaptic architecture components suggest a widespread disruption likely impacting neurotransmission. Glutamate stands as the principal excitatory neurotransmitter; any disturbances in its receptor interaction can tip the scales in neuronal excitability, synaptic adaptability, and general cerebral activity [58]. When juxtaposed with a diminished GABA receptor function-key in moderating inhibitory neurotransmission-this scenario underscores a concerning landscape of synaptic anomalies [59].

The KEGG pathway analysis illuminated a connection between the diminished gene profiles and established neurodegenerative pathways, notably including PD, Huntington’s, and ALS. Of these, PD stands out, mainly due to the degenerative impact on DA neurons in the substantia nigra. A growing body of research postulates a central role for mitochondrial dysfunction in PD’s pathogenesis [60]. Our data strengthens this hypothesis, spotlighting the mitochondrial anomalies linked to POLG in patient-derived DA neurons. In essence, these insights suggest that mitochondrial impairments, magnified by *POLG* mutations, could serve as a foundational nexus for various neuronal malfunctions, possibly predisposing individuals to neurodegenerative ailments, with PD at the forefront. These mitochondrial perturbations, combined with potential synaptic challenges, collectively undermine neuronal well-being, laying down the groundwork for future therapeutic exploration and disease control approaches.

In synthesizing of the pronounced upregulation of pathways such as Apoptosis, ErbB signaling, Cell cycle, Mitophagy (animal), Regulation of actin cytoskeleton, and the Wnt signaling pathway, it’s evident that the ramifications of *POLG* mutations extend far beyond mere mitochondrial disturbances, also trigger a ripple effect across a broad spectrum of cellular activities, especially in DA neurons. These cascading dysfunctions, rooted in mitochondrial anomalies and extending to neuronal signaling and structural pathways, craft a multifaceted landscape of neuronal compromise. This complex web of disturbances, dominated by POLG-associated mitochondrial dysregulation, sets the stage for heightened vulnerability to neurodegenerative conditions. As such, these revelations not only deepen our understanding of POLG-linked pathologies but also illuminate potential avenues for targeted therapeutic measures, fostering hope for innovative interventions in the battle against neurodegenerative diseases.

## Conclusion

In our extensive research, we meticulously produced and profiled MOs from pluripotent stem cells, unraveling the intricacies of midbrain neuron differentiation and their possible ties to neurodegenerative conditions. One pivotal discovery was the stark contrast observed between control and POLG mutant MOs. The POLG variant MOs distinctly presented with pronounced mitochondrial anomalies, as well as notable neuroanomalies, all underpinned by our GO analysis. Detailed analysis revealed a downregulated gene profile in POLG patient-derived DA neurons, indicating possible disruptions in mitochondrial pathways, notably the oxidative phosphorylation pathways, as well as neurological challenges related to neurotransmission and synaptic functions.

Interestingly, we observed that POLG MOs had about a 50% reduction in DA neuron levels, i.e., 17% as compared to 30% in controls. Moreover, the single-cell transcriptomic assessment of these MOs unveiled significant disparities in their profiles vis-à-vis the controls. A recurring theme throughout our findings was the interplay between mitochondrial respiratory chain complexes and the oxidative phosphorylation pathway, pinpointing it as a potential hub in neurodegenerative mechanisms. Additionally, our KEGG analysis showcased the predisposition of POLG-derived DA neurons towards well-established neurodegenerative pathways.

In summation, our data firmly establishes the crucial impact of *POLG* mutations on midbrain neuron well-being, particularly highlighting mitochondrial functionality and synaptic coherence. This investigation not only enriches our grasp of the intricate scenarios tied to neurodegenerative maladies related to *POLG* mutations but also paves the path for specialized therapeutic interventions moving forward.

## Data Availability

The RNA sequencing analysis read count data can be accessed in NCBI Gene Expression Omnibus (GEO) data deposit system with an accession number GSE241743. All other data are available from the corresponding author upon request.

## Author’s contributions

K.L contribute to the conceptualization; A.C and T. Y contribute to the methodology; the investigation; K.L and A.C contribute to the writing original draft; all authors contribute to writing review and editing; K.L contribute to the funding acquisition and to the resources; K.L contributes to the supervision. All authors agree to the authorships.

## Acknowledgements

We thank members of the Molecular Imaging Centre and Flow Cytometry Core Facility for their expertise and assistance in confocal imaging and flow cytometry data recording.

---

**Table 1:** List of cell numbers in scRNA-seq.

| Sample | DA | Total no. |
| --- | --- | --- |
| Control | 5353 | 17698 |
| Patient | 743 | 4871 |

**Figure S1.**
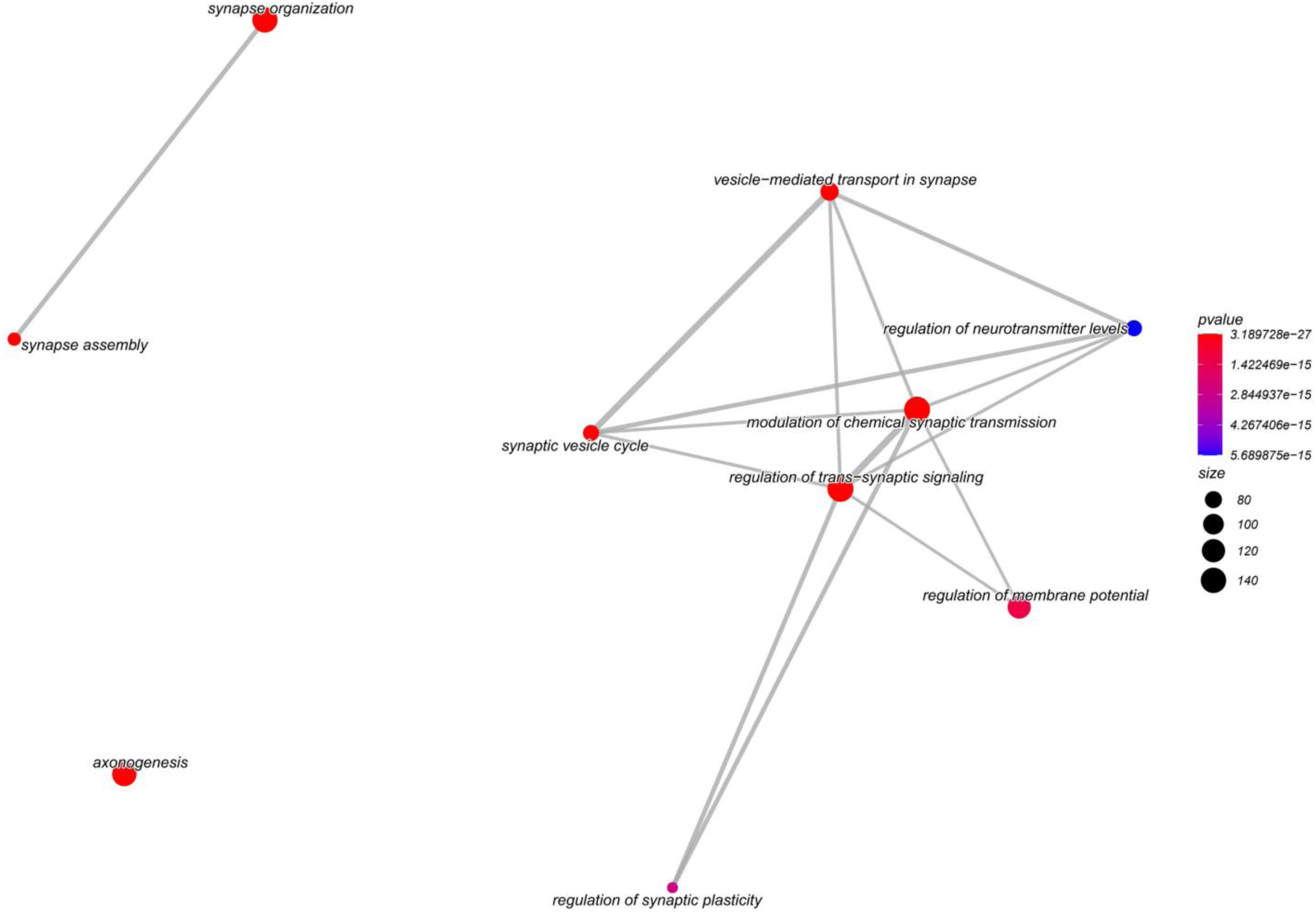
GO biological process emapplot analysis of downregulated DEGs in hMOs of POLG patients compared with controls.

**Figure S2.**
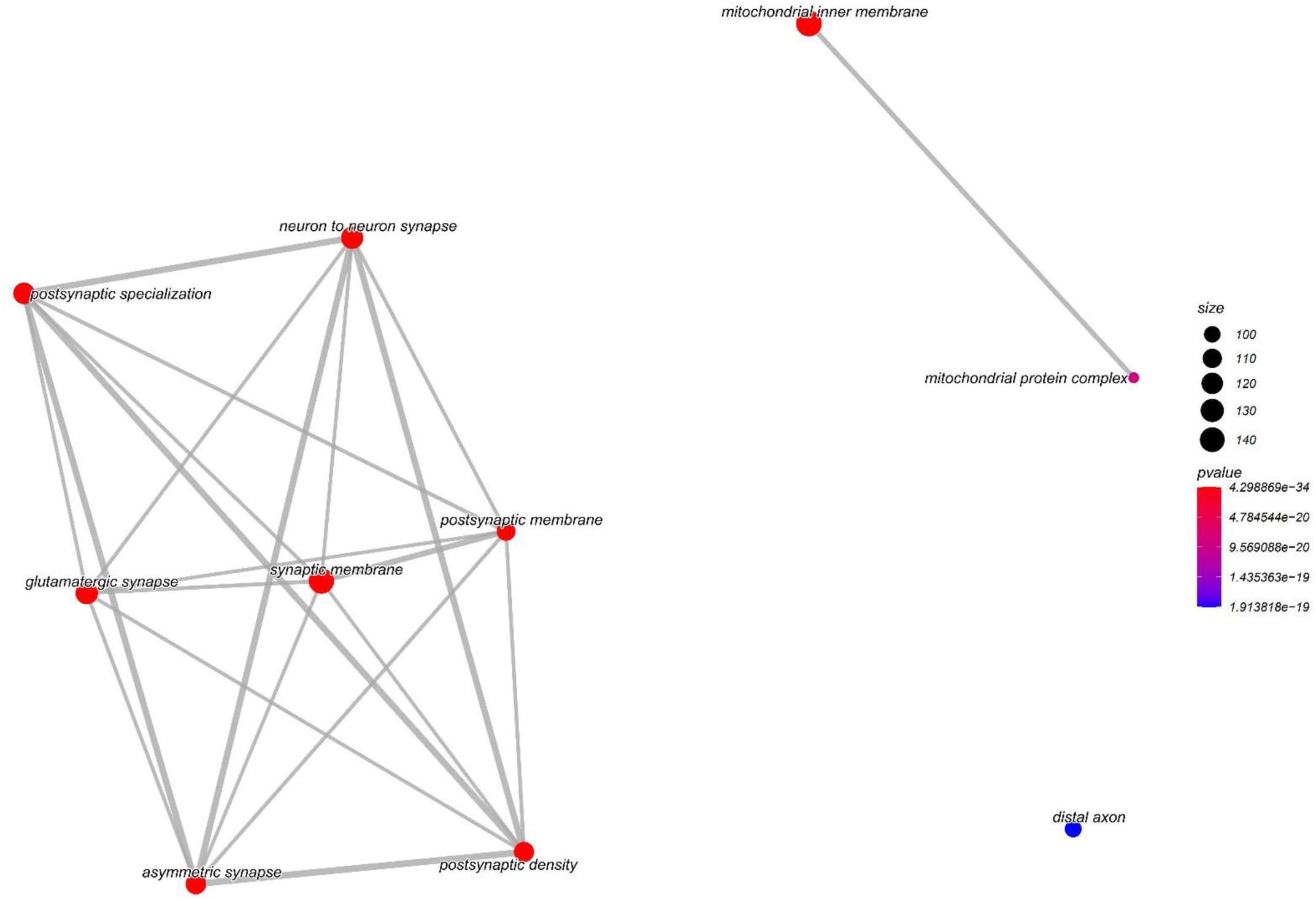
GO cell component emapplot analysis of downregulated DEGs in hMOs of POLG patients compared with controls.

**Figure S3.**
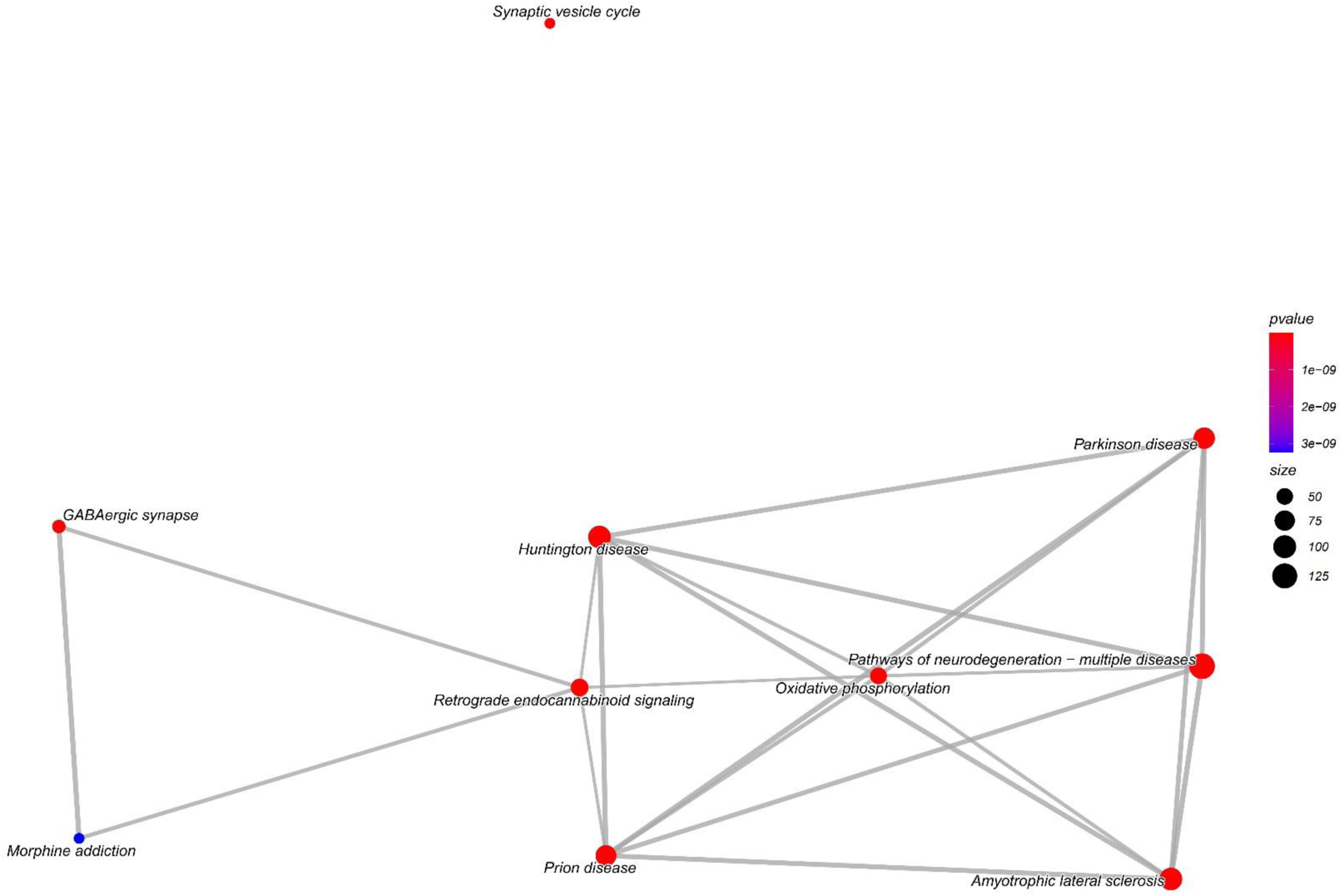
KEGG pathway emapplot analysis of downregulated DEGs in hMOs of POLG patients compared with controls.

**Figure S4.**
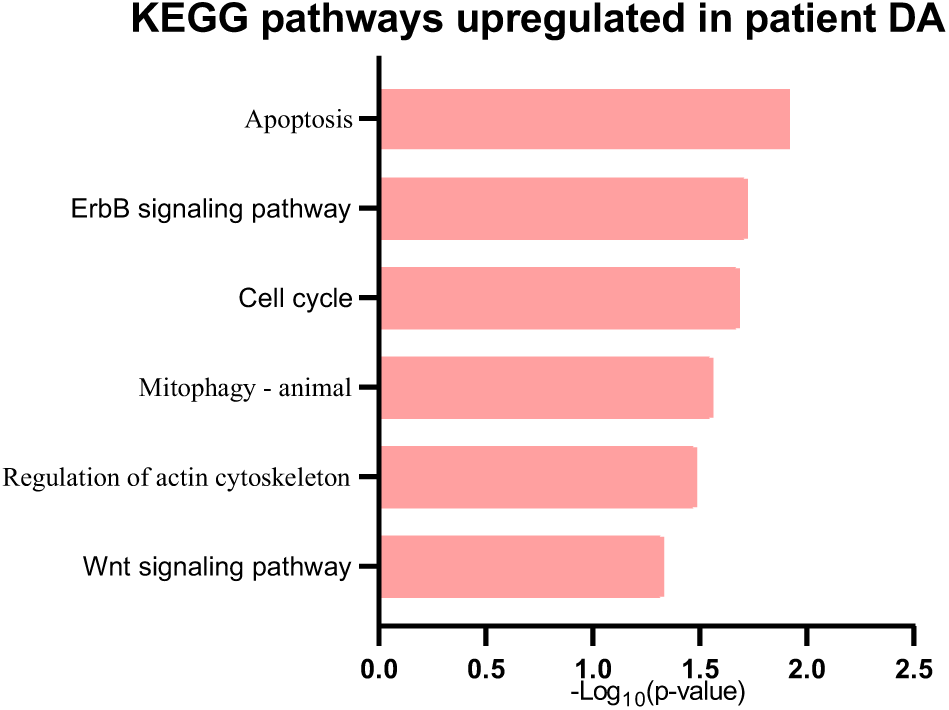
KEGG pathway analysis of upregulated DEGs in DA of hMOs of POLG patients compared with controls.

**Table S1.**
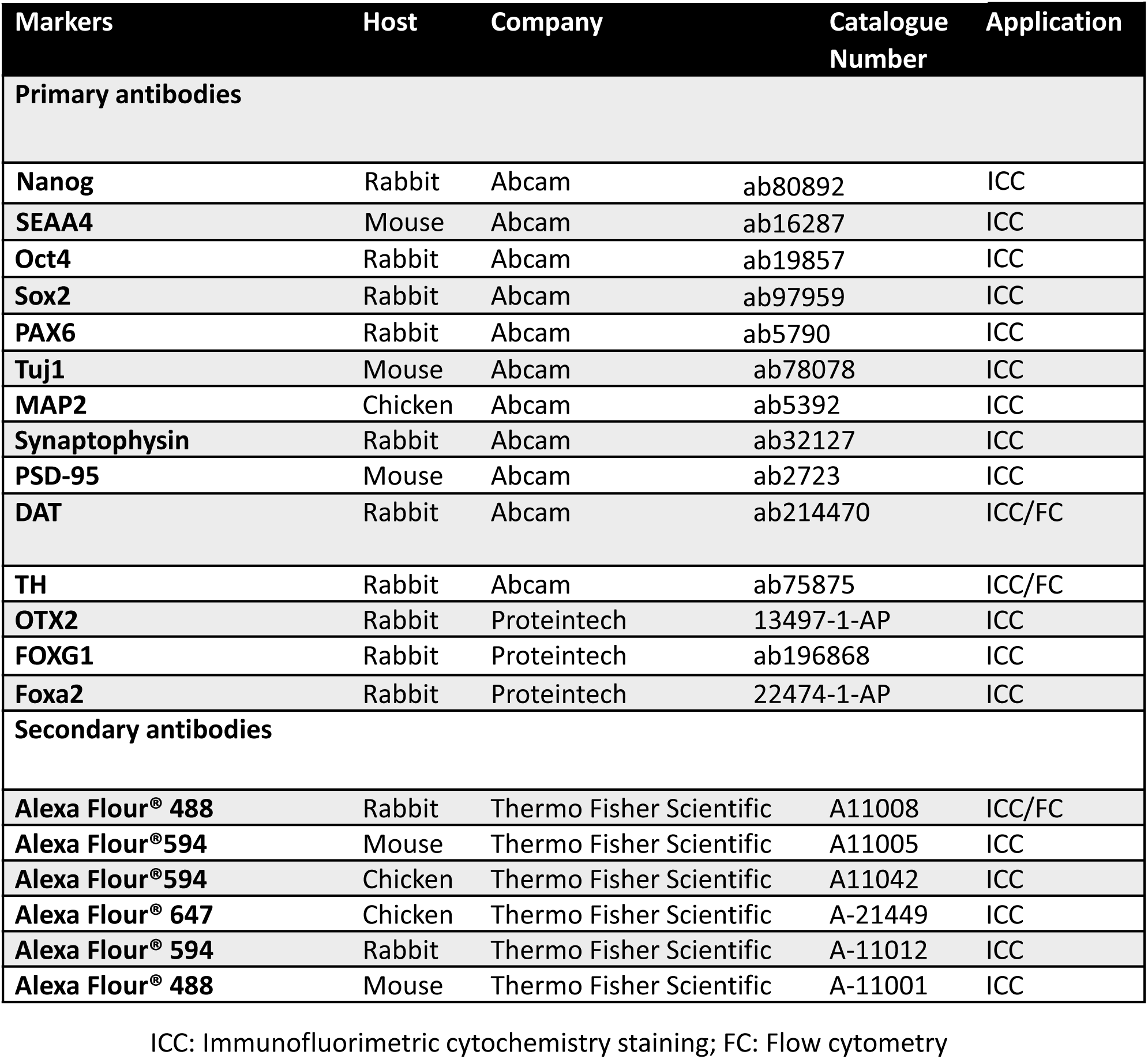
List of antibodies used in the study.

**Table S2.**
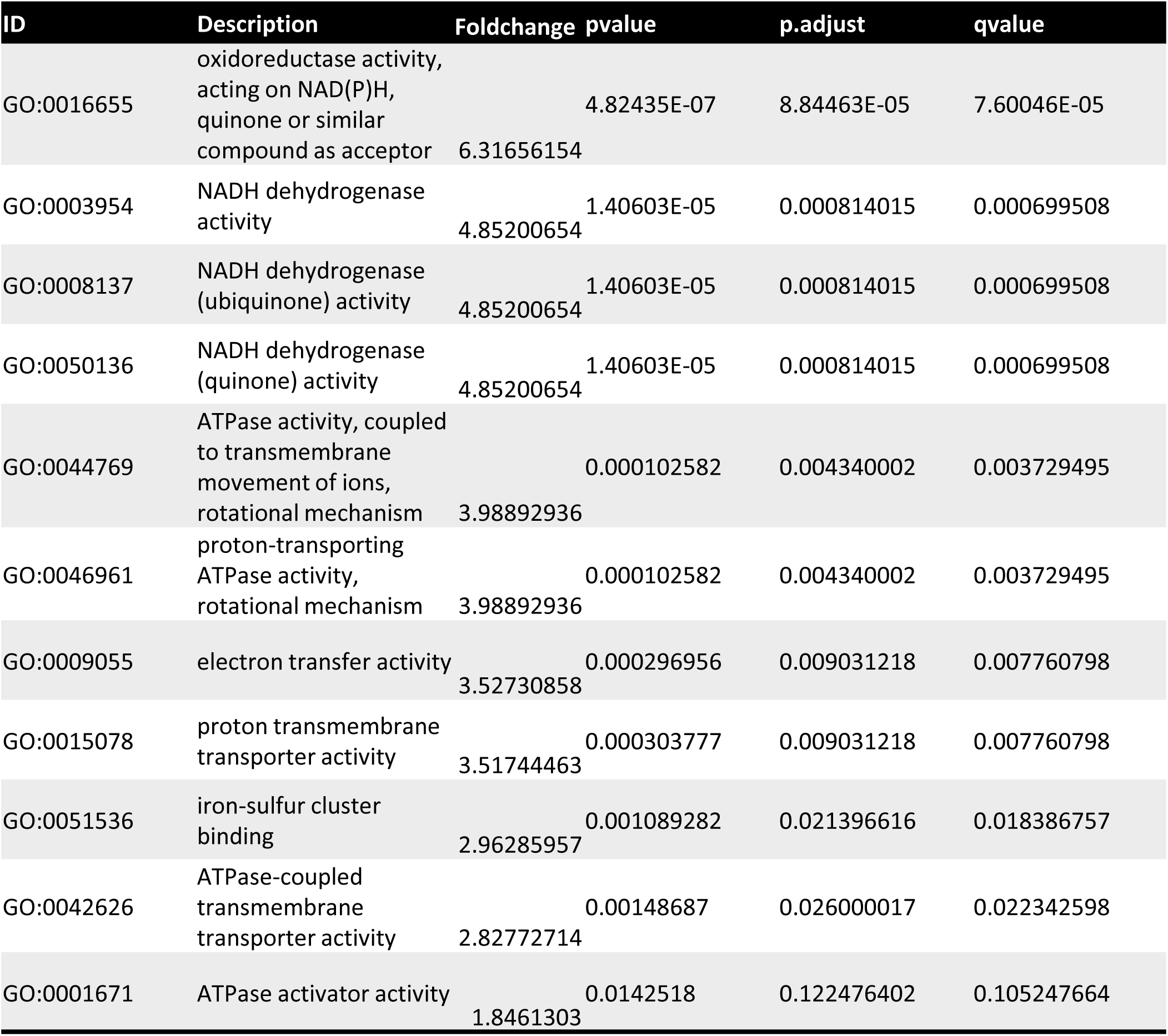
Mitochondrial related GO/MF downregulated in patient DA.

| ID | Description | Foldchange | pvalue | p.adjust | qvalue |
| --- | --- | --- | --- | --- | --- |
| GO:0016655 | oxidoreductase activity, acting on NAD(P)H, quinone or similar compound as acceptor | 6.31656154 | 4.82435E-07 | 8.84463E-05 | 7.60046E-05 |
| GO:0003954 | NADH dehydrogenase activity | 4.85200654 | 1.40603E-05 | 0.000814015 | 0.000699508 |
| GO:0008137 | NADH dehydrogenase (ubiquinone) activity | 4.85200654 | 1.40603E-05 | 0.000814015 | 0.000699508 |
| GO:0050136 | NADH dehydrogenase (quinone) activity | 4.85200654 | 1.40603E-05 | 0.000814015 | 0.000699508 |
| GO:0044769 | ATPase activity, coupled to transmembrane movement of ions, rotational mechanism | 3.98892936 | 0.000102582 | 0.004340002 | 0.003729495 |
| GO:0046961 | proton-transporting ATPase activity, rotational mechanism | 3.98892936 | 0.000102582 | 0.004340002 | 0.003729495 |
| GO:0009055 | electron transfer activity | 3.52730858 | 0.000296956 | 0.009031218 | 0.007760798 |
| GO:0015078 | proton transmembrane transporter activity | 3.51744463 | 0.000303777 | 0.009031218 | 0.007760798 |
| GO:0051536 | iron-sulfur cluster binding | 2.96285957 | 0.001089282 | 0.021396616 | 0.018386757 |
| GO:0042626 | ATPase-coupled transmembrane transporter activity | 2.82772714 | 0.00148687 | 0.026000017 | 0.022342598 |
| GO:0001671 | ATPase activator activity | 1.8461303 | 0.0142518 | 0.122476402 | 0.105247664 |

**Table S3.** Neural related GO/MF downregulated in patient DA.

| ID | Description | Foldchange | pvalue | p.adjust | qvalue |
| --- | --- | --- | --- | --- | --- |
| GO:0035254 | glutamate receptor binding | 9.30086037 | 5.00195E-10 | 5.50215E-07 | 4.72816E-07 |
| GO:0044325 | ion channel binding | 7.81422956 | 1.53381E-08 | 8.43593E-06 | 7.24925E-06 |
| GO:0030276 | clathrin binding | 7.28343369 | 5.20674E-08 | 1.31065E-05 | 1.12628E-05 |
| GO:0035255 | ionotropic glutamate receptor binding | 4.59338485 | 2.55044E-05 | 0.001402742 | 0.001205419 |
| GO:0051082 | unfolded protein binding | 4.56431693 | 2.72699E-05 | 0.001428422 | 0.001227486 |
| GO:0008017 | microtubule binding | 3.64798731 | 0.000224912 | 0.008510334 | 0.007313187 |
| GO:0098918 | structural constituent of synapse | 3.63704095 | 0.000230653 | 0.008510334 | 0.007313187 |
| GO:0000149 | SNARE binding | 3.61536355 | 0.000242458 | 0.008603347 | 0.007393115 |
| GO:0043014 | alpha-tubulin binding | 3.57466123 | 0.00026628 | 0.008973183 | 0.007710926 |
| GO:0030165 | PDZ domain binding | 3.55765592 | 0.000276913 | 0.008973183 | 0.007710926 |
| GO:0016917 | GABA receptor activity | 3.55696725 | 0.000277353 | 0.008973183 | 0.007710926 |

